# The Effects of Histone H2B ubiquitylations and H3K79me_3_ on Transcription Elongation

**DOI:** 10.1101/2023.01.05.522859

**Authors:** Mai T. Huynh, Bhaswati Sengupta, Wladyslaw A. Krajewski, Tae-Hee Lee

## Abstract

Post-translational modifications of histone proteins often mediate gene regulation by altering the global and local stability of the nucleosome, the basic gene-packing unit of eukaryotes. We employed semi-synthetic approaches to introduce histone H2B ubiquitylations at K34 (H2BK34ub) and K120 (H2BK120ub) and H3 K79 trimethylation (H3K79me3). With these modified histones, we investigated their effects on the kinetics of transcription elongation by RNA Polymerase II (Pol II) using single-molecule FRET. Pol II pauses at several locations within the nucleosome for a few seconds to minutes, which governs the overall transcription efficiency. We found that H2B ubiquitylations suppress pauses and shorten the pause durations near the nucleosome entry while H3K79me3 shortens the pause durations and increases the rate of RNA elongation near the center of the nucleosome. We also found that H2BK34ub facilitates partial rewrapping of the nucleosome upon Pol II passage. These observations suggest that H2B ubiquitylations promote transcription elongation and help maintain the chromatin structure by inducing and stabilizing nucleosome intermediates and that H3K79me3 facilitates Pol II progression possibly by destabilizing the local structure of the nucleosome. Our results provide the mechanisms of how these modifications coupled by a network of regulatory proteins facilitate transcription in two different regions of the nucleosome and help maintain the chromatin structure during active transcription.

## INTRODUCTION

The nucleosome is the most basic gene packaging unit in eukaryotes that contains ∼147 base-pair (bp) DNA wrapping around a histone protein core made of two H2A-H2B dimers and one (H3-H4)_2_ tetramer ^1,2^. Histone proteins are targets for various post-translational modifications (PTMs) such as acetylation, methylation, phosphorylation, and ubiquitylation ^3-6^. Many histone PTMs play gene regulatory roles by mediating the interactions among the nucleosome, transcription factors, chromatin remodelers, and other various enzymes or by altering the global and local structure and stability of the nucleosome ^7-9^. Lysine methylations and ubiquitylations can have either activating or repressive roles depending on their locations ^10-15^. Ubiquitylations of H2B at K34 and K120 have been associated with enhanced processivity of RNA Polymerase II (Pol II) ^16^. These two modifications are coupled in the same network of protein interactions where the MOF-MSL and RNF/20/40 complexes ubiquitylate H2B K34 and K120, respectively, in an inter-dependent manner through interactions with a general transcription elongation factor Paf1C ^16^. An *in vitro* study has found that H2BK34 ubiquitylation (H2BK34ub) destabilizes H2A-H2B dimer association with the nucleosome, thus facilitating hexasome formation by histone chaperone Nap1 ^17-19^. Recent Cryo-EM studies of H2BK34 nucleosomes reported that the ubiquitin moieties protrude the DNA gyre gap in the nucleosome and distort the DNA near the nucleosome entry region, thereby destabilizing the nucleosome and facilitating the binding of a histone methyltransferase Dot1L ^20,21^. However, no report is currently available to show how such effects of H2BK34ub directly regulate the kinetics of transcription elongation by Pol II. An important feature of the transcription kinetics through the nucleosome is that Pol II faces numerous kinetic barriers. These barriers are imposed by strong DNA-histone interactions which induce pauses of Pol II. These pauses can last for a few seconds to minutes and thus govern the overall efficiency of transcription since the actual elongation rate of RNA is very fast at 40 – 50 nucleotides per second ^22,23^. K120 is another target for ubiquitylation at H2B. While ubiquitylation at H2BK120 (H2BK120ub) exhibits only a moderate impact on nucleosome stability, it has been studied more actively for its role in recruiting Dot1L to methylate histone H3 at K79, which has been coupled to active transcription ^17,24-32^. Trimethylation at H3 K79 (H3K79me_3_) promotes transcription *in vitro* and is associated with highly active genes ^32-34^ although there has not been any report showing how H3K79me_3_ alters the transcription kinetics through the nucleosome. A structural study hinted at a possible role for dimethylated H3K79 (H3K79me_2_) inducing a rearrangement of the local histone surface while no change was observed in the global structure of the nucleosome ^35^.

Here, we investigated the effects of H2BK34ub, H2BK120ub, H3K79me_3_, and H3K79me_3_/H2BK120ub on the kinetics of transcription elongation by Pol II through the nucleosome and the status of DNA rewrapping after Pol II passage. We employed semi-synthetic approaches to introduce these modifications and a highly refined *in vitro* transcription system where we used single-molecule FRET (smFRET) to monitor the nucleosome dynamics during and after Pol II passage in real-time in a time-resolved manner. Our approaches and system enabled investigations of transcription kinetics at the nucleosome level without any interference from unknown or ambiguous factors, which will help interpret the previous and future *in vivo* results in depth. Our results indicate that H2B ubiquitylations and H3K79me_3_ accelerate transcription via two different mechanisms. H2B ubiquitylations facilitate transcription by suppressing pauses at the nucleosome entry and shortening the entry pause durations while H3K79me_3_ shortens the pause durations and increases the rate of elongation toward the internal region of the nucleosome. We also found that H2BK34ub results in a significantly increased population of nucleosome intermediates with partially rewrapped DNA upon Pol II passage. These results provide the mechanisms of how H2BK34ub, H2BK120ub, and H3K79me_3_ that are coupled by a network of regulatory proteins facilitate transcription in two different regions of the nucleosome and how H2BK34ub may help keep the structural integrity of chromatin during active transcription.

## MATERIAL AND METHODS

### Histone Preparation

Human histone proteins H3, H4, and H3 C110A K79C were purchased from the Histone Source (Colorado State University, Fort Collins, CO). Plasmids containing wild type human histone proteins H2A and H2B were kindly provided by Dr. Song Tan (Pennsylvania State University, University Park, PA). Plasmids expressing human H2B mutants with either K34C or K120C mutation were generated via site-directed mutagenesis. Histone H3 with K79 trimethylation was prepared following published protocols ^36^. Briefly, histone protein H3 with K79C mutation was dissolved in Alkylation buffer (1M HEPES, 10 mM D/L-methionine, 4 M Guanidium-HCl, 20 mM DTT) and incubated for 1 hour at 37 °C. Then 100 mg of (2-bromoethyl) trimethylammonium bromide was added to the reaction mixture, followed by heating at 50 °C with occasional stirring for 2.5 hours. An aliquot of 10 μL 1 M DTT was then added, and the reaction was allowed to proceed for another 2.5 hours. Excess β-mercaptoethanol (BME) was added to the reaction mixture to quench the reaction. The reaction results in the product trimethylated aminoethylcysteine which has the β carbon of the trimethylated lysine side chain replaced with a sulfur (i.e. the methyl group linkage replaced with a thioether linkage). This mimetic has been shown to reproduce biochemical activities of H3K79me_3_^36^. We will call this mimetic H3K79me_3_ for simplicity. The product was purified using PD-10 columns (GE Healthcare, Chicago, IL) in deionized water containing 3 mM BME. The product was then lyophilized and stored at –80 °C. The product was analyzed with mass spectrometry to confirm its modification and purity (Fig. S1). Histone H2B with K34 or K120 ubiquitylation was prepared following published protocols ^37,38^. A plasmid expressing His-TEV-ubiquitin G76C was acquired from Addgene (Watertown, MA) and the protein was purified using Talon beads (Takara Bio Inc., Japan) following published protocols ^37,38^. Briefly, ubiquitin and histone H2B were both dissolved in 50 mM Tris-HCl (pH 8.6) and 6M Urea. They were then mixed with the molar ratio of two histone proteins per one ubiquitin followed by the addition of TCEP to 5 mM. The reaction mixture was incubated at room temperature for 30 minutes. Dichloroacetone was then added, and the reaction was allowed to proceed for 16 hours on ice. The reaction mixture was quenched by 10 mM BME. The reaction results in a nonhydrolyzable ubiquitylated histone H2B mimetic which has been used for biochemical and structural studies of H2B ubiquitylated nucleosomes ^37,38^. We call these H2B ubiquitylation mimetics H2BK34ub and H2BK120ub for simplicity. The product was purified using HisPur NiNTA column with 10 μL TEV protease at 4 °C to cleave 6xHis tag. SDS-PAGE was used to confirm the product (Fig. S2). The product was then dialyzed against deionized water, lyophilized, and stored at –80 °C.

### DNA Preparation

Nine DNA fragments (see Table. S1 for sequences) were purchased (Integrated DNA Technologies, Coralville, IA). The fragments were annealed and ligated to form nucleosomal DNA containing the Widom 601 sequence ^39^. The EC-42 strands (see Table. S1 for sequences) with a 9-nt mismatch followed by a G-less cassette were ligated to the nucleosome entry site. The +34 and +112 nucleotides counted from the nucleosome entry site were labeled with NHS-ester functionalized Atto647N and Cy3 via the amine-functionalized C6 linkers coupled to the thymine bases (Integrated DNA Technologies, Coralville, IA). Another FRET pair was used where the +57 and +138 nucleotides counted from the nucleosome entry site were labeled with the same FRET pair, NHS-ester functionalized Atto647N and Cy3.

### Nucleosome Reconstitution

Histone proteins were mixed in stoichiometric amounts and eluted through a HiLoad 16/600 Superdex 200pg column (GE Healthcare) to produce H2A-H2B dimers and (H3-H4)_2_ tetramers separately ^40^. Nucleosomes were reconstituted by dialyzing a stoichiometric mixture of histones and DNA in a dialysis device (Slide-A-Lyzer MINI Dialysis Device, 7K MWCO, Thermo Fisher Scientific) against 1X TE (pH 8.0) buffer with stepwise dilution of salt at 850, 650, 500, 300, and 2.5 mM NaCl ^40,41^. In total, five sets of nucleosomes were assembled: nucleosomes with no modification (wild type), H2BK34ub, H2BK120ub, H3K79me_3_ and H3K79me_3_/H2BK120Kub. All assembled transcription templates containing the nucleosome were gel-purified by the crush and soak method and confirmed by native PAGE (Fig. S3).

### Pol II and TFIIS Purification

Yeast Pol II was purified from *Saccharomyces Cerevisiae* as described in a previous publication ^42^. Briefly, yeast cells expressing Pol II with a TAP tag at Rpb4 were grown in YPD media with 20 μg/mL adenine sulfate. Yeast cells were harvested by centrifugation at 6,000 g and stored at – 80 °C in Lysis buffer (250 mM Tris-HCl pH 7.5, 5 mM EDTA, 50 μM ZnCl_2_, 50 % glycerol, 5 % DMSO, 4 mM leupeptin, 5 mM pepstatin A, 2 mM aprotinin, 2 mM benzamidine, 10 mM DTT). Yeast cells were then thawed, lysed by homogenizer (EmulsiFlex-C3, AVESTIN, Canada), and centrifuged at 6000 g to collect the supernatant. The supernatant was centrifuged at 20,000 rpm twice for an hour and filtered through a sterilized gauze pad to remove any lipid residue. The filtered supernatant was purified with an IgG-Sepharose column followed by purification with a MonoQ column at 4 °C. The elution fractions were collected and analyzed by SDS-PAGE to confirm the presence and purity of yeast Pol II.

A plasmid to express His-tagged TFIIS was kindly provided by Dr. Joseph Reese (Pennsylvania State University, University Park, PA). His-tagged TFIIS was expressed in *BL21(DE3)pLysS* cells in 2-YT media with 100 μg/mL ampicillin. Cells were harvested by centrifugation at 6,000 g and stored at –80 °C in Lysis buffer (20 mM Tris-HCl pH 7.5, 0.5 M NaCl, 10 mM imidazole, 10 μM ZnCl_2_, 10 % glycerol, 1 mM PMSF, 2 mM benzamidine). Cells were then thawed, lysed by homogenizer (EmulsiFlex-C3, AVESTIN, Canada), and centrifuged at 16,000 g for 30 min to collect the supernatant. The supernatant was incubated with Talon beads (Takara Bio Inc., Japan) for an hour at 4 °C. The beads were then washed multiple times and eluted with 150 mM imidazole. The elution was collected in 0.5 mL fractions and analyzed by SDS-PAGE.

### Formation of Transcription Elongation Complex

The assembled transcription template was diluted in 50 μL transcription buffer (60 mM KCl, 1 mM MnCl_2_, 50 mM K-HEPES (pH 7.8), 0.5 mM DTT, and 10 % glycerol) to 10 nM followed by addition of 9.5 nM Pol II, 0.3 mM UpG primer, 0.1 mg/mL BSA, and 2 unit/μL RNasin® RNase inhibitor. The reaction mixture was incubated for 5 min at 30 °C to allow Pol II docking on the 9-nt DNA bubble. To initiate transcription, a mixture of ATP, CTP, and UTP each at 1mM was added to the Pol II-DNA complex, and then incubated for another 30 min at 30 °C. Due to the lack of GTP in the system, the elongation process is halted at the end of the G-less cassette.

### Single-Molecule FRET Measurements

#### Surface Preparation for smFRET Measurements

Pre-drilled microscope quartz slides were purchased from G. Finkenbeiner Inc. (Waltham, MA). The slides were cleaned thoroughly following the published protocols ^43^. Briefly, the slides were immersed in acetone, dichloromethane, and methanol in series and sonicated for 15 min each. They were then subjected to KOH etching and finally dipped in a Nochromix® (Alconox, White Plains, NY) solution in concentrated sulfuric acid. Finally, they were dried using nitrogen, then coated with 1:50 biotin-PEG-silane (Laysan Bio Inc., Arab, AL) and PEG-silane (Laysan Bio Inc., Arab, AL). Five flow channels were constructed on the slide.

#### Nucleosome Dynamics during Transcription Elongation by Pol II

A streptavidin solution (40 μL, 0.1 mg/mL) was injected into a channel constructed on a microscope slide and incubated for 15 mins for biotin-streptavidin conjugation. After 15 minutes, unbound streptavidin was removed by washing with 40 μL wash buffer (10 mM NaCl and 10 mM K-HEPES, pH 7.8). A diluted transcription mixture (∼1 nM final nucleosome concentration) was injected into the channel and incubated for 5 minutes for the nucleosomes to conjugate to streptavidin on the surface. After 5 minutes the unbound nucleosomes were washed with wash buffer containing protocatechuic acid (10 mM, MilliporeSigma, MA) and protocatechuate 3,4-dioxygenase (1 unit/uL, MilliporeSigma, MA). The slide was then placed on an smFRET microscope which is custom-built on a commercial inverted microscope (Nikon TE2000, Japan). Right before recording the fluorescence images, the imaging buffer (a complete set of NTP at 0.4 mM, 30 nM TFIIS, 1 unit/mL PCD, 10 mM PCA, and 1 mM Trolox in the transcription buffer, 30 μl) was injected into the channel. The fluorescence images were collected every 200 ms integrating 200 ms fluorescence signal on two spectral channels to separate the FRET acceptor signal (Atto647N, 635 nm) from that of the donor (Cy3, 532 nm) using an EMCCD (Ixon Ultra, Andor Technology, Ireland). Each stack of fluorescence images forms a movie file that was recorded for 3 minutes. To make sure that any change in the FRET efficiency was not due to Atto647N photo-bleaching, we switched on a red laser (632nm, CrystaLaser, Reno NV) to excite the acceptor directly at the end of each movie, and only those points with an acceptor signal were subject to further analysis. A total of 5 movies (15 minutes total) were collected from each channel. The time traces of fluorescence intensities from each pair of Cy3 and Atto647N were extracted from the fluorescence images.

#### Nucleosome unwrapping and rewrapping during and after Transcription by Pol II

The transcription reaction mixture was injected into a channel and incubated for 5 min at room temperature. The imaging buffer was then added to wash unbound nucleosomes and resume the transcription process. Immediately after the addition of the imaging buffer, three consecutive fluorescence images were recorded every 4.5 sec with 532 nm laser excitation followed by another three images with 635 nm laser excitation. All images were taken with 150 ms signal integration.

The signal from 635 nm excitation is used to verify no photobleaching or blinking of Atto647N. Each stack of fluorescence images that forms a movie file was taken for 3 minutes. Under our excitation conditions, the vast majority of Atto647N was not photobleached or blinking during imaging ^44^. Ten movies from different spots were recorded and analyzed. For each of all five sets of nucleosomes, at least 5 independent measurements were made.

### Transcription Efficiency Assay

The transcription elongation complex was generated as mentioned above except Cy3-UTP was used in the NTP mix to resume transcription at the end of the G-less cassette. The transcription reaction were incubated for 30 minutes at 30 °C and subsequently terminated by 2X stop buffer (30 mM Tris-HCl pH 8.0, 100 mM NaCl, 5 mM EDTA, and 1% SDS). The resulting transcription reaction was incubated with Proteinase K for 30 min at 37 °C to release RNA from the elongation complex, and then with streptavidin agarose beads for 1 hour to pull down biotinylated DNA. RNA was extracted with PCIAA (Phenol:Chloroform:Isoamyl alcohol) and precipitated with chilled ethanol and 4μg glycogen. Solid RNA was then dissolved in RNA loading buffer (5 mM EDTA in formamide), heated at 65°C for at least 10 min, and loaded to 8% urea gel. Fluorescent gel images were taken with Typhoon (GE Healthcare) and analyzed with ImageJ for intensities.

## RESULTS

### A single-molecule FRET system reports nucleosome dynamics during transcription

Previous studies have established that Pol II pauses at several DNA-histone contact points during transcription through the nucleosome ^45-49^. Using smFRET, we investigated the pause frequency, durations, and the elongation rate between pauses to reveal the effects of H2BK34ub, H2BK120ub, and H3K79me_3_ (Fig. 1A). The fluorophores were labeled at the 34^th^ and 112^th^ nucleotides of the non-template strand of the nucleosomal DNA counted from the entry site, reporting DNA unwrapping and rewrapping during and after the Pol II passage. The FRET pair locations are designed to give a high-FRET efficiency when DNA is properly wrapped and to show stepwise FRET decreases as DNA unwraps during Pol II passage (Fig. 1B). Without Pol II or NTP, no such FRET changes are observed for 15 minutes. We monitored the changes in FRET during Pol II passage revealing three stable states with visually distinct FRET efficiencies (Fig. 1B). These stable FRET states correspond to the paused states of the elongation complex at various locations in the nucleosome (Fig. S4 A-D).

**Figure 1.**
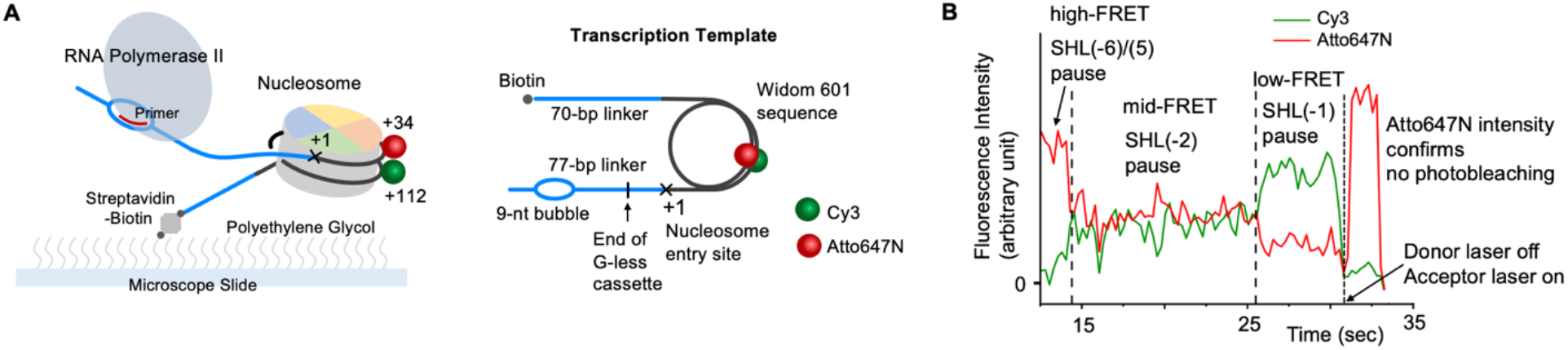
An smFRET experimental setup shows detailed nucleosome dynamics during transcription elongation by Pol II. (A) The structure of the transcription template containing a 601 nucleosome sequence and surface immobilization scheme to monitor smFRET signals from the nucleosome. A FRET pair (Cy3 and Atto647N for donor and acceptor, respectively) was labeled at the nucleosomal DNA, reporting the nucleosome dynamics during Pol II progression. (B) An example showing the intensity traces of the FRET donor (Cy3, green) and the acceptor (Atto647N, red) during Pol II progression through the nucleosome. A high-FRET state indicates a pause event at SHL(−6) or SHL(−5), a mid-FRET state indicates a pause event at SHL(−2), and a low-FRET state indicates a pause event at SHL(−1) according to the structural analysis with the FRET pair locations (Fig. S4).

We named the three stable FRET states high-, mid-, and low-FRET state (Fig. 1B). We assigned the high-FRET state to the early pauses at superhelical location (SHL) (−6) and (−5), the mid-FRET to the mid pauses at SHL (−2), and the low-FRET to the late pauses at SHL (−1) according to the estimated FRET efficiencies from the nucleosome structures with paused Pol II (Fig. S4 A-D)^46^. Of note, only a small number of traces show the mid-FRET state, which is consistent with a previously published report that SHL(−2) is a minor swift pause ^46^.

### Ubiquitylated nucleosomes suppress early pauses and shorten the pause duration

We counted the number of nucleosomes showing the early pauses at SHL(−6) and SHL(−5) in the first 2 minutes after the NTP addition for Pol II progression. The early pause frequency was measured by dividing the number of nucleosomes starting with a high-FRET state by the number of total Cy3 fluorescence spots. The Cy3 spots include nucleosomes that have not been initiated for transcription, are being transcribed, and have been done with transcription. Assuming that the initiation efficiency is constant across all templates, the fraction of nucleosomes showing high FRET represents the pause density or frequency. Our results indicate that the pause frequency in the cases of H2K34ub, H2BK120ub, and H3K79me_3_/H2BK120ub nucleosomes is lower than that in the unmodified case (Fig. 2). These results suggest that H2K34ub and H2BK120ub facilitate transcription by suppressing early pauses. Of note, the extent of the effect is a lot more significant with H2K34ub.

**Figure 2.**
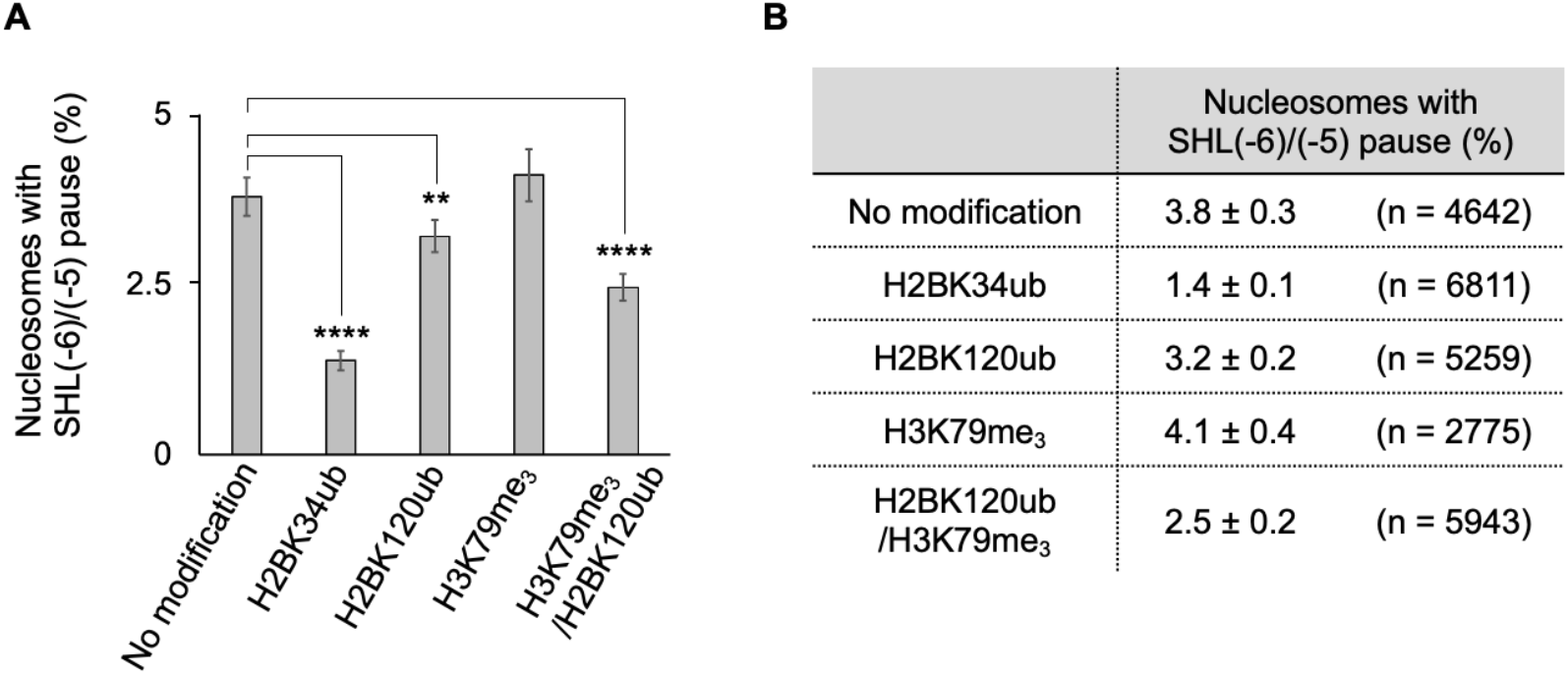
H2B ubiquitylations show a significantly reduced frequency of pauses at SHL(−6)/(−5). (A) The fractions of nucleosomes showing a high-FRET state out of all nucleosomes immobilized on the surface reveal that H2B ubiquitylations suppress early pauses at SHL(−6)/(−5). (B) Numerical data for the histogram shown in A. The errors are the standard errors for binomial distributions with the given sample sizes. The binomial distribution *p-*values are <0.0001, 0.010, 0.82, and <0.0001 for H2BK34ub, H2BK120ub, H3K79me_3_, and H2BK120ub/H3K79me_3_, respectively. The number of asterisks designates the level of significance in the difference.

Next, we measured the pause durations which are the durations of the high-FRET states. The FRET duration histograms fit well to a single exponential decay whose time constant is the average pause duration (Fig. 3). Note that the histogram starts at 30 sec time point which is typically the time spent on reactant injection, flow-channel mounting, and microscope focusing. The high-FRET pause duration corresponding to the SHL(−6)/(−5) pause duration is significantly shorter in the cases of the ubiquitylated nucleosomes (13.7 ± 1.8, 16.7 ± 1.6, and 16.8 ± 1.9 sec respectively for H2BK34ub, H2BK120ub, H3K79me_3_/H2BK120ub) than in the cases of the unmodified (22.8 ± 1.9 sec) or H3K79me_3_ nucleosomes (22.5 ± 2.5 sec) (Fig. 3). It has been reported based on *in vivo* and *in vitro* studies that ubiquitylated nucleosomes promote Pol II processivity and facilitate hexasome generation by Nap1 ^16,17^. As hexasome generation by Nap1 takes place near the DNA-(H2A-H2B) contact region, H2B ubiquitylations may destabilize the nucleosome in the entry region where SHL(−6)/(−5) are located. Therefore, our results suggest that the facilitated transcription upon H2B ubiquitylation is due to suppressed pauses and shortened pause durations in the SHL(−6)/(−5) region.

**Figure 3.**
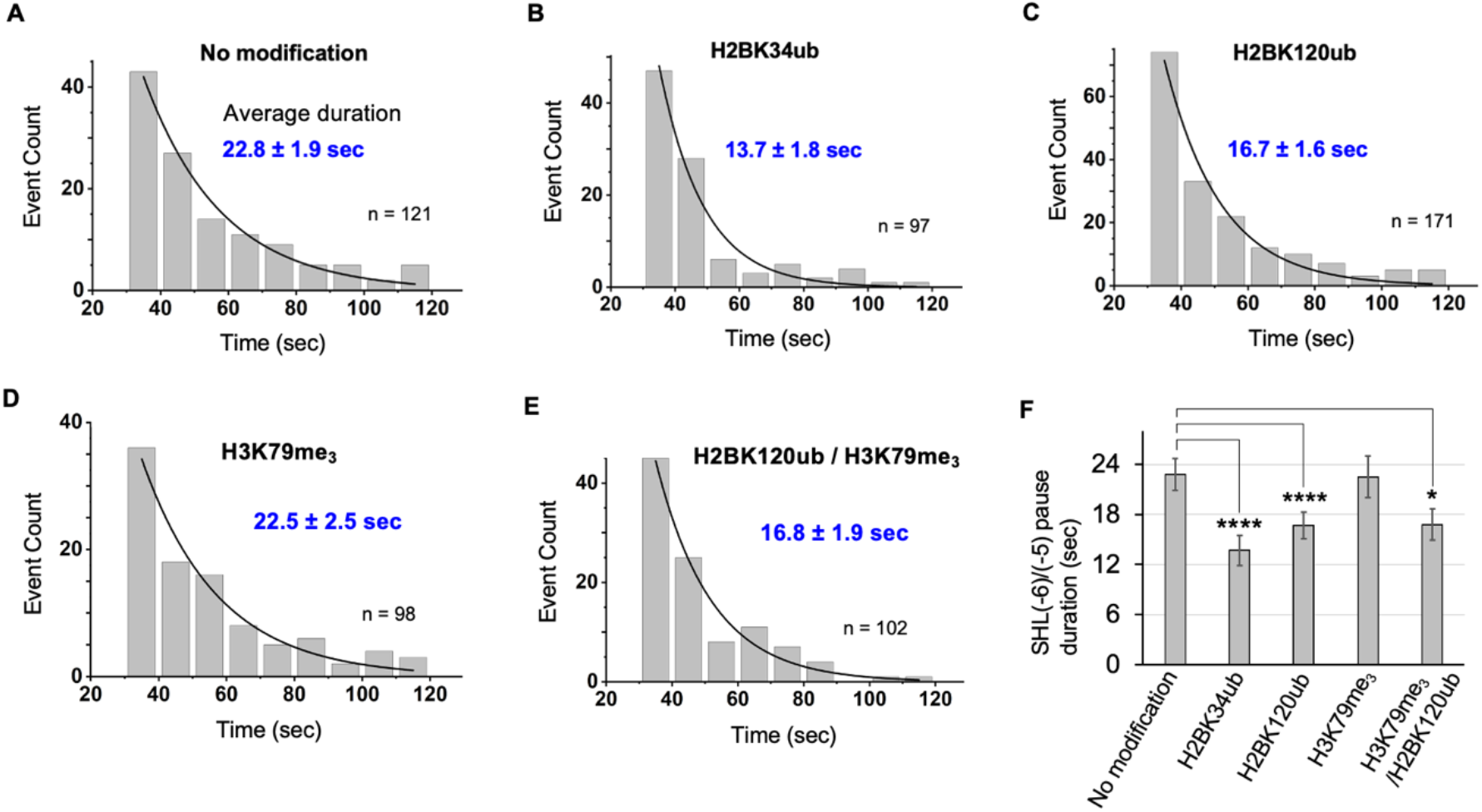
Histone H2B ubiquitylations shorten the duration of the early pauses at SHL(−6) and SHL(−5). The high-FRET state duration or lifetime histograms are shown for (A) unmodified, (B) H2BK34ub, (C) H2BK120ub, (D) H3K79me_3_, and (E) H2BK120ub/H3K79me_3_ nucleosomes. The sample size n for each case is shown in each chart. The average and error values are from the fitting with an exponential decay function. (F) Summary of the pause durations showing shortened pause durations by H2BK34ub and K120ub. The one-sided exponential distribution *p*-values (power = 0.8) are <0.0001, <0.0001, 0.760 and 0.015 for H2BK34ub, H2BK120ub, H3K79me_3_, and H2BK120ub/H3K79me_3_, respectively. The number of asterisks designates the level of significance in the shortened pause duration.

### Nucleosomes with H3K79me_3_ show increased transcription rates and reduced pause durations in the SHL(−2)/(−1) region

To investigate the effects of H3K79me_3_ on transcription, we measured the elongation times between pauses. We identified FRET transitions from a high-to a low-FRET state between which elongation takes place (Fig. 4A, Fig. S5). The transition time windows include pauses at SHL(−2) which are often short-lived (Fig. S5). The histograms of the width of the transition time windows are well fit with a single exponential decay function (Fig. 4B-F). According to the results, the H3K79me_3_ nucleosomes show a faster elongation rate (0.83 ± 0.06 and 0.70 ± 0.04 sec or 48 ± 3 and 57 ± 3 nucleotides per second (nt/sec), respectively for H3K79me_3_ and H3K79me_3_/H2BK120ub nucleosomes) than the other nucleosomes (1.15 ± 0.12, 1.01 ± 0.17, and 1.04 ± 0.15 sec or 35 ± 4, 40 ± 7, and 38 ± 6 nt/sec, respectively for unmodified, H2BK34ub, and H2BK120ub nucleosomes) (Fig. 4B-F). To estimate the rates in nt/sec, we assume that the early pauses are mostly at SHL(−5) as pauses at SHL(−6) are much weaker ^46,50,51^. With the low-FRET pauses at SHL(−1), the elongation time is for transcribing 40 nucleotides. The results suggest that H3K79me_3_ facilitates transcription by elevating the elongation rate from the entry to the SHL(−1) region. As no change in the global nucleosome structure has been identified in the H3K79me_2_ nucleosome structure ^35^, we suggest that this effect may be due to local changes induced by H3K79me_3_ that would be similar to those by H3K79me_2_. Such changes may affect the conformational flexibility of the nucleosome near this region, which could weaken the already swift SHL(−2) pauses and facilitate Pol II passage. This suggestion led to a hypothesis that H3K79me_3_ may shorten SHL(−2)/(−1) pause durations. To test this hypothesis, we moved the FRET pair locations near SHL(−2) at the 57^th^ and 138^th^ nucleotides and repeated smFRET measurements for the unmodified and the H3K79me_3_ nucleosomes (Fig. 5A). We observed two stable FRET states – high- and low-FRET states (Fig. 5B). All pauses from SHL(−6) to SHL(−1) should show a high-FRET state according to the structural analysis (Fig. S4 E-H). As the elongation rate is very fast compared to the pause durations, the moment of a transition from a high- to a low-FRET state should be the moment when Pol II passes through SHL(−1) at the observation time resolution of 200 ms. Accordingly, the high-FRET duration should be a convolution of all pause durations from SHL(−6) to SHL(−1). A change in the high-FRET duration should indicate a change in the pause duration at SHL(−2)/(−1) since H3K79me_3_ does not induce any change in the pause durations at SHL(−6)/(−5). The distribution of the high-FRET duration clearly shows significantly shorter pauses for the H3K79me_3_ nucleosomes (Fig. 5D, 47.2 sec on average) than the unmodified nucleosomes (Fig. 5C, 56.3 sec on average), validating our hypothesis on the shortened pause durations at SHL(−2)/(−1) by H3K79me_3_. These results show that H3K79me_3_ facilitates Pol II passage through the nucleosome by shortening the durations of pauses at SHL(−2)/(−1), thereby elevating the rate of elongation up to SHL (−1).

**Figure 4.**
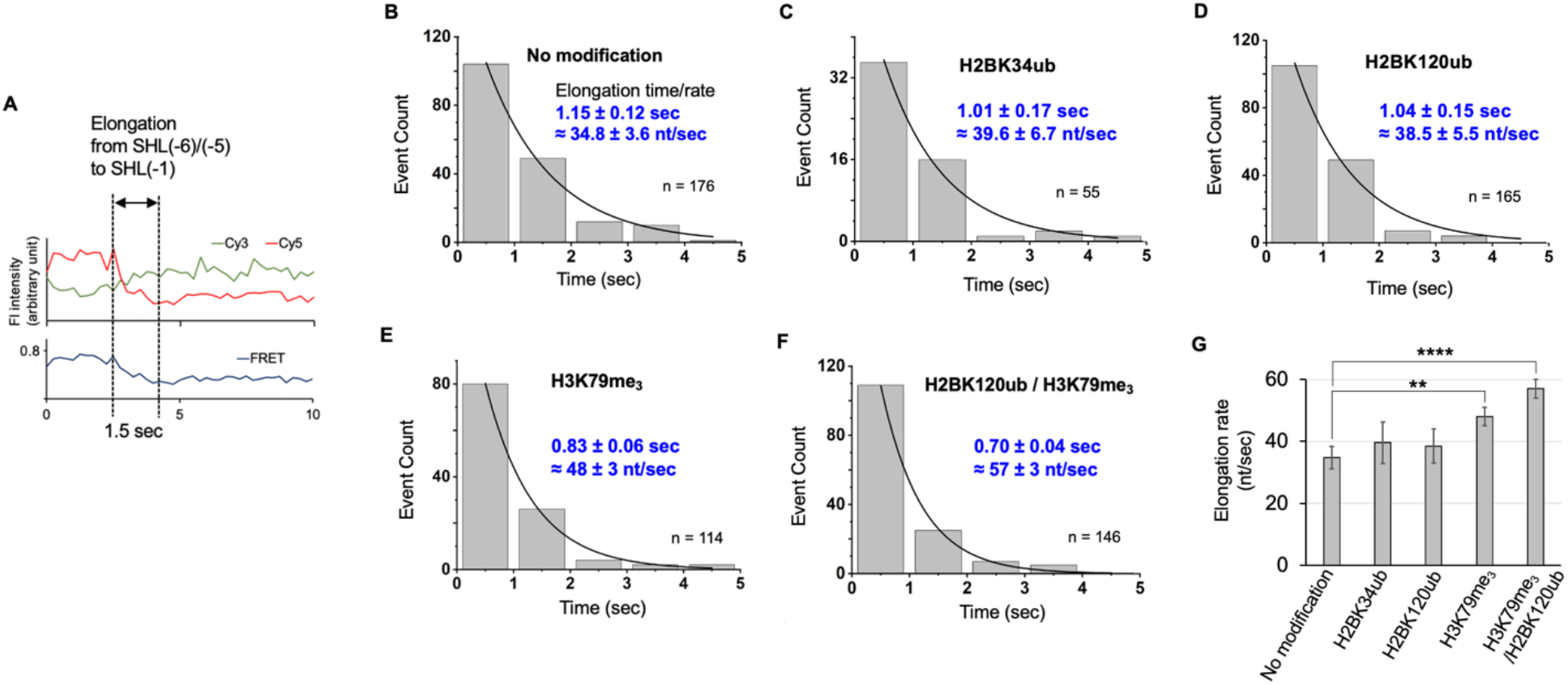
H3K79me_3_ increases the rate of transcription elongation. (A) An example of fluorescence intensity traces demonstrating the elongation phase of transcription from SHL(−6)/(−5) to SHL(−1) that is indicated by a FRET transition from a high- to a low-FRET state. Elongation time histograms are shown for (B) unmodified, (C) H2BK34ub, (D) H2BK120ub, (E) H3K79me_3_, and (F) H2BK120ub/H3K79me_3_ nucleosomes. The sample size n for each case is shown in each chart. The elongation time and error are from fitting each histogram with an exponential decay function. The elongation rate is estimated by assuming that early pauses are mostly at SHL(−5) and consequently the measured time is to elongate 40 nucleotides. (G) Summary of the results show that H3K79me_3_ increases the elongation rate. The one-sided exponential distribution *p*-values (power = 0.8) are 0.447, 0.324, 0.006, and <0.0001 for H2BK34ub, H2BK120ub, H3K79me_3_, and H2BK120ub/H3K79me_3_, respectively. The number of asterisks designates the level of significance in the increase.

**Figure 5.**
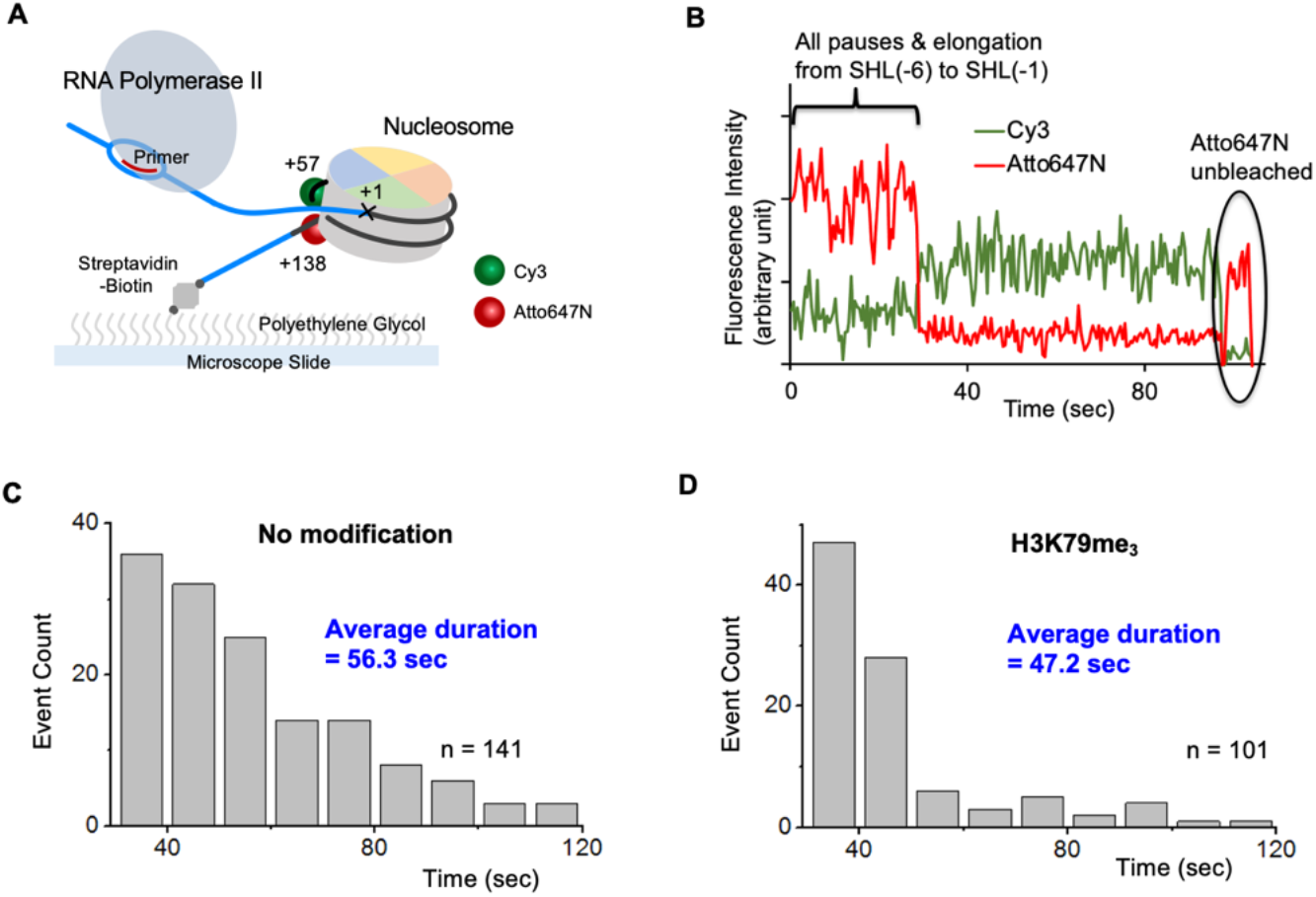
H3K79me_3_ shortens late pauses at SHL(−2)/(−1). (A) Experimental setup with new FRET pair locations to investigate the late pause dynamics at SHL(−2)/(−1). According to the structural analysis with the FRET locations (Fig. S4), a high-FRET state is expected until Pol II passes through SHL(−1). (B) An example of fluorescence intensity traces shows an abrupt transition from a high- to a low-FRET state indicating Pol II progression through SHL(−1). High-FRET state duration histograms are shown for (C) unmodified and (D) H3K79me_3_ modified nucleosomes. The average durations were calculated directly from the durations and their counts. These durations represent the convoluted durations of pauses from SHL(−6) to SHL(−1). A significantly shorter duration is shown in D, indicating that H3K79me_3_ shortens the late pause durations at SHL(−2)/(1) as no change was observed in the early pause durations (Fig. 3).

### *In vitro* transcription assay confirms facilitated transcription

We performed an *in vitro* transcription assay where fluorescent RNA products from transcription for 30 min were extracted and analyzed on a urea gel in all five cases of the nucleosomes (Fig. 6A). Three different levels of signal integration time were used to visualize the three different regions of the gel (Fig. 6B). Higher amounts of RNA products are evident in all of the modified nucleosome cases, confirming facilitated transcription whose mechanisms were revealed by the smFRET measurements (Fig. 6B-C). The change is insignificant in the case of H2BK120ub as the effect was only weak in the single-molecule measurements as well. The assay also showed a considerably large amount of RNA products from H3K79me_3_/H2BK120ub nucleosomes as the result should be the sum of the effects from H3K79me_3_ which affects the SHL(−2)/(−1) region and H2BK120ub which affects the SHL(−6)/(−5) region.

**Figure 6.**
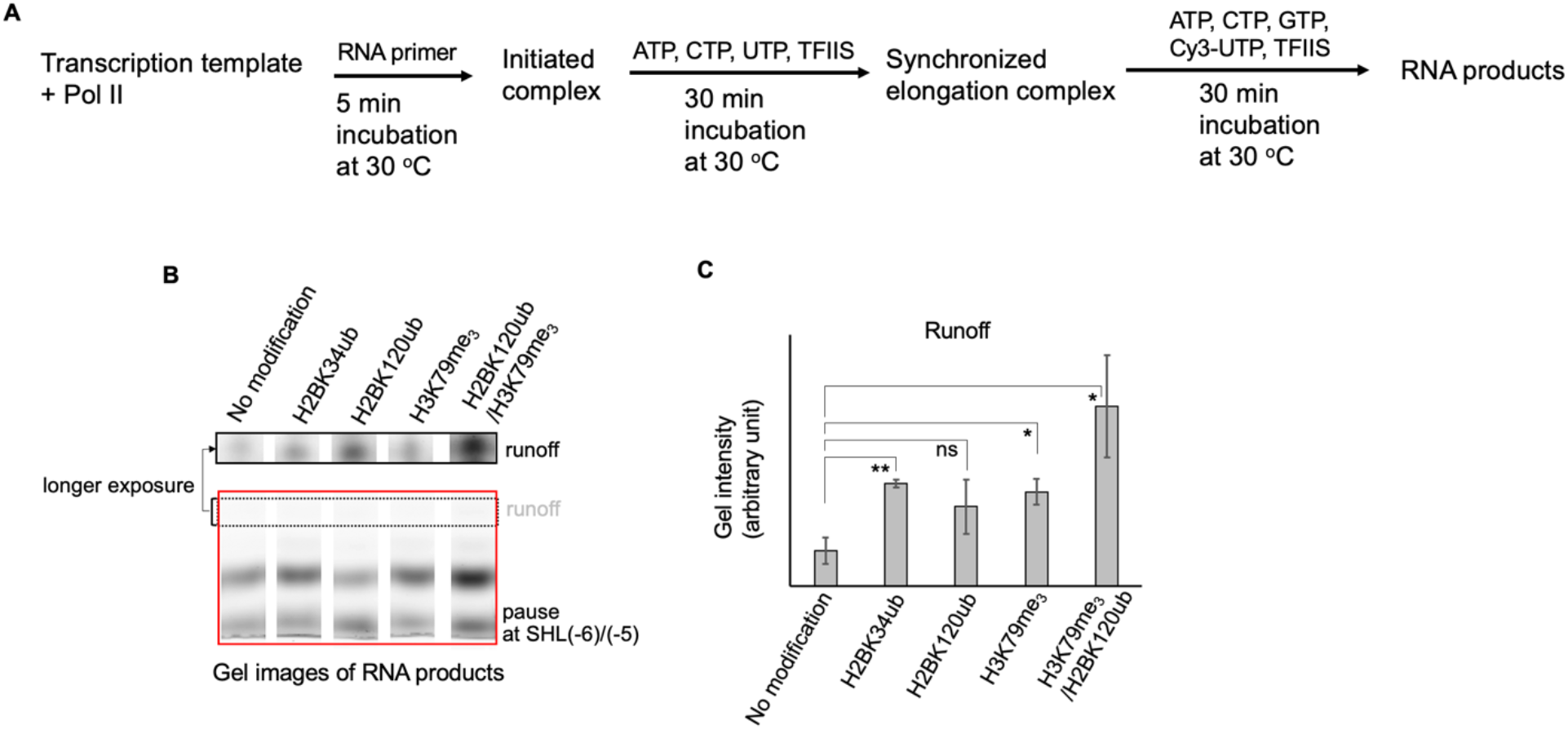
An *in vitro* transcription assay confirms that H2B ubiquitylations and H3K79me3 increases transcription efficiency. (A) The experimental scheme for the *in vitro* transcription assay utilizing fluorescent UTP labeled with Cy3. (B) Gel images of the RNA product incorporated by Cy3-UTP. The entire gel image is shown in the red box. Four bands are apparent. The lowest band should contain the RNA products up to the earliest pauses at SHL(−6)/(−5) as the synchronized elongation complex at the end of the G-less cassette contains no fluorescent nucleotides. The highest band should represent the runoff product. Two different signal integration/exposure times were used to image the gels more clearly in the runoff region. The black dotted-box region shows the longest RNA product on the full gel (red box). The black dotted-box region with a longer exposure time is shown in the black solid-box (runoff). (C) The amounts of the runoff products measured from the intensities of the runoff bands shown in B indicate significantly increased efficiency of transcription elongation in all of the modified nucleosomes except for the H2BK120ub case. The error bars are standard deviations from duplicate measurements. The one-sided *p-*values in the cases of H2BK34ub, H2BK120ub, H3K79me_3_, and H2BK120ub/H3K79me_3_, are 0.010, 0.089, 0.023, and 0.030, respectively. The number of asterisks designates the level of significance in the increased RNA production and “ns” stands for “not significant”.

### H2K34ub facilitates partial DNA rewrapping upon Pol II passage

Nucleosome recovery after transcription is vital to maintaining the chromatin structure for proper gene regulation. Here, we investigated the fate of the nucleosome in the entry proximal DNA-(H2A-H2B) region after Pol II passage with the FRET pair depicted in figure 1A. We modified the smFRET measurement scheme to avoid premature photobleaching of fluorophores and ensure that a FRET change is due to nucleosome dynamics rather than Atto647N blinking. In the modified scheme, fluorophores were imaged for a cycle of 1.05 sec every 4.5 sec for a total of 3 min. Each imaging cycle is made of a 450 ms period of donor excitation at 532 nm followed by a 150 ms dark period and another 450 ms period of weak acceptor excitation at 635 nm (Fig. 7A). The two signals from 532 nm excitation (i.e. Cy3 signal and Atto647N signal via FRET) and the signal from 635 nm excitation (i.e. Atto647N signal from direct excitation) are plotted together at a single time point (Fig. 7BC). We excluded traces that show a sign of Atto647N photobleaching or blinking (i.e. extinction of Atto647N signal from the direct 635 nm excitation cycle) from further analysis. A FRET transition from a high- to a lower FRET state indicates DNA unwrapping (Fig. 7BC). Upon Pol II passage, nucleosomes may rewrap DNA completely to a high-FRET state (Fig. 7B). We counted the number of nucleosomes showing the signature of complete and stable nucleosome rewrapping after unwrapping and calculated the fraction of such nucleosomes (Fig. 7D). The fractions are 14.9 ± 3.6, 19.7 ± 4.0, 18.2 ± 3.9, 14.4 ± 3.5, and 19.3 ± 3.9 % for the unmodified, H2BK34ub, H2BK120ub, H3K79me_3_, and H3K79me_3_/H2BK120ub nucleosomes, respectively, indicating no significant difference. In addition to the signature of complete DNA rewrapping, we also observed a FRET transition from a low- to a higher FRET state which does not recover fully to the high-FRET level where DNA unwrapping started (Fig. 7C). This state indicates DNA rewrapping to an intermediate state, which we call partial rewrapping of DNA. We counted such nucleosomes and calculated the fractions (Fig. 7E). The fractions are 14.3 ± 3.5, 36.7 ± 4.0, 16.8 ± 4.8, 20.2 ± 3.7, and 24.0 ± 4.3 % for the unmodified, H2BK34ub, H2BK120ub, H3K79me_3_, and H3K79me_3_/H2BK120ub nucleosomes, respectively, indicating a significant increase in partial DNA rewrapping by H2BK34ub. The sum of the full and partial DNA rewrapping fractions are 29.2 ± 5.0, 56.4 ± 6.0, 35.0 ± 5.3, 34.6 ± 5.3, and 43.2 ± 5.8 % for the unmodified, H2BK34ub, H2BK120ub, H3K79me_3_, and H3K79me_3_/H2BK120ub nucleosomes, respectively, indicating that more than half of H2BK34ub nucleosomes have partially or fully rewrapped DNA upon Pol II passage. Structural studies have suggested that ubiquitin may have stable nonspecific interactions with DNA and may also interact with histone proteins via its acidic regions ^20,52,53^. Therefore, the interactions between the ubiquitin and the nucleosome can help the nucleosome retain the ubiquitylated H2A-H2B dimer within the complex, which may help reassembly of the nucleosome. As H2BK120ub nucleosomes displayed no significantly increased rewrapping, we suggest that this effect depends on the location of ubiquitin. As the two ubiquitin molecules in H2BK34ub nucleosomes may protrude between the two gyres of the nucleosomal DNA, they may contact DNA readily to assist H2A-H2B retention and DNA rewrapping while the ubiquitin molecules in H2BK120ub nucleosomes are farther away from histone-DNA interfaces to interact effectively with DNA.

**Figure 7.**
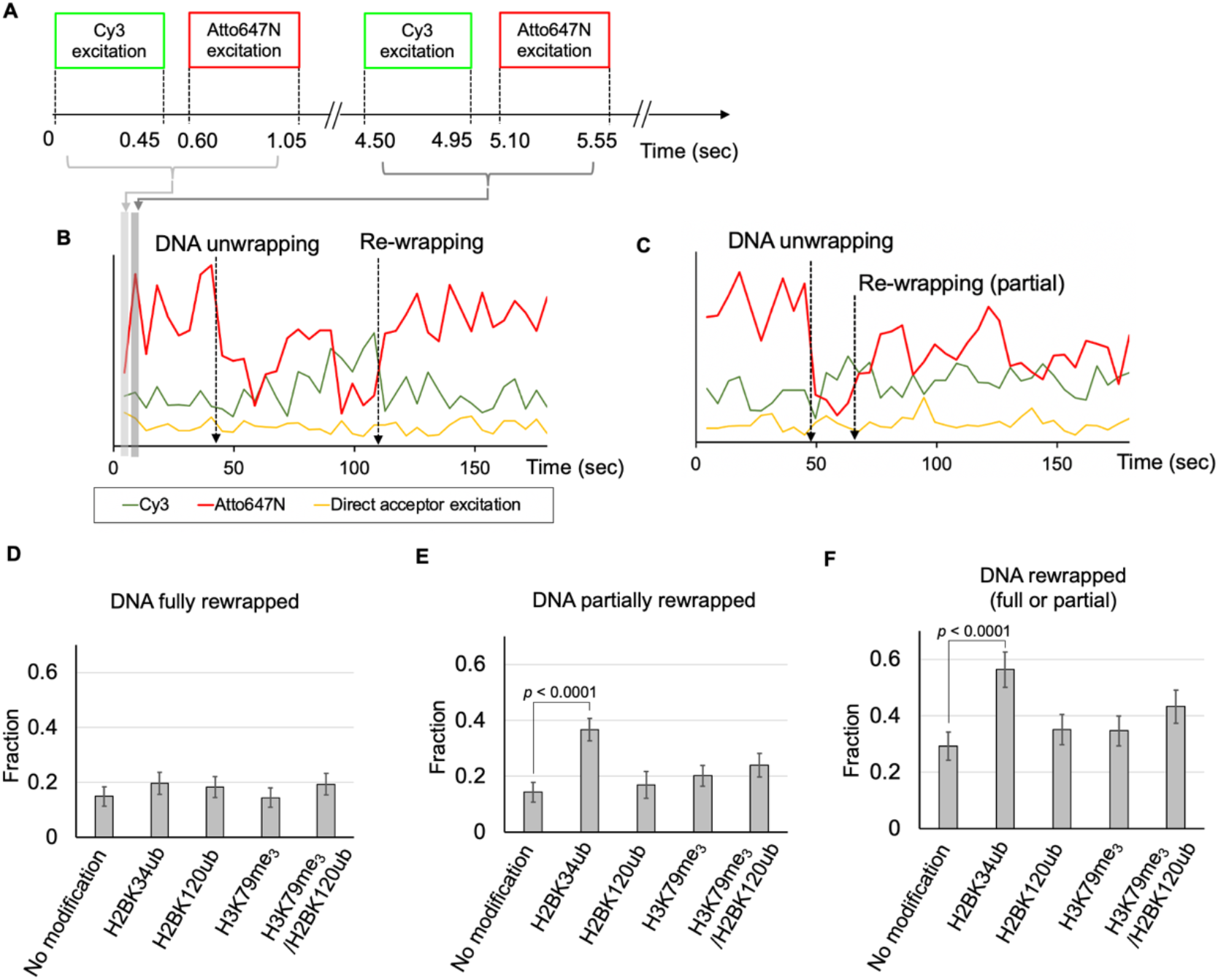
H2BK34ub facilitates partial DNA rewrapping upon Pol II passage. (A) The intermittent FRET excitation scheme to investigate the status of the nucleosomal DNA unwrapping and rewrapping without photobleaching of the fluorophores and interference from Atto647N blinking for 3 minutes. (B) An example of fluorescence intensity traces shows the signatures of DNA unwrapping and rewrapping. A high-FRET state in the beginning indicates the intact nucleosome before transcription. Some FRET dynamics follow as Pol II progresses through the nucleosome. The complex reaches a high-FRET state at around ∼110 sec time point, suggesting complete DNA rewrapping. (C) An example of fluorescence intensity traces shows the signatures of DNA unwrapping and partial rewrapping. Upon Pol II progression coinciding with some FRET dynamics in the ∼50 – 65 sec window, the complex reaches a state with fluctuations between mid- to high-FRET, which is not the completely rewrapped state, suggesting partial DNA rewrapping. (D) The fractions of the nucleosomes showing the signature of DNA rewrapping are constant among all nucleosomes investigated. The fractions are 14.9 ± 3.6, 19.7 ± 4.0, 18.2 ± 3.9, 14.4 ± 3.5, and 19.3 ± 3.9 % for the unmodified, H2BK34ub, H2BK120ub, H3K79me3, and H3K79me3/H2BK120ub nucleosomes, respectively. (E) The fraction of the nucleosomes showing the signature of partial DNA rewrapping is significantly higher with H2BK34ub. The fractions are 14.3 ± 3.5, 36.7 ± 4.0, 16.8 ± 4.8, 20.2 ± 3.7, and 24.0 ± 4.3 % for the unmodified, H2BK34ub, H2BK120ub, H3K79me3, and H3K79me3/H2BK120ub nucleosomes, respectively. The one-sided binomial distribution *p*-value for a higher fraction with H2BK34ub is <0.0001. (F) The sum of the fractions in D and E show the fractions of the nucleosomes with full or partial DNA rewrapping upon Pol II passage. The fractions are 29.2 ± 5.0, 56.4 ± 6.0, 35.0 ± 5.3, 34.6 ± 5.3, and 43.2 ± 5.8 % for the unmodified, H2BK34ub, H2BK120ub, H3K79me3, and H3K79me3/H2BK120ub nucleosomes, respectively. The sample sizes are 167, 259, 214, 105, and 192 nucleosomes that do not show any sign of blinking or photobleaching for 3 minutes after transcription start for the unmodified, H2BK34ub, H2BK120ub, H3K79me3, and H3K79me3/H2BK120ub nucleosomes, respectively. The one-sided binomial distribution *p*-value for a higher fraction with H2BK34ub is <0.0001. All the errors shown in D-F are the standard errors for binomial distributions with the given sample sizes.

## DISCUSSIONS

Histone proteins are rich targets for various PTMs that are often associated with gene regulation. Some mimetics of these PTMs have been implemented with semi-synthetic methods to construct highly-refined *in vitro* experimental systems. For example, histone methylations via aminoethylation of cystine have been published to successfully reproduce their biochemical functions ^35,54,55^. A disulfide coupling approach for ubiquitylation has also been published to reproduce the functions of ubiquitylated histones ^21,27,38^. We employed these approaches and a highly refined *in vitro* transcription system to construct a tractable single-molecule transcription system to investigate the effects of histone H2B ubiquitylations and H3K79me_3_ on the kinetics of transcription elongation through the nucleosome and the status of DNA wrapping during and after transcription ^43^.

Pol II pauses at various locations within the nucleosome during transcription ^46,50,51^. Out of these pause locations, we focus on the first four - SHL(−6), (−5), (−2), and (−1) among which SHL(- 5) and (−1) induce major pauses while SHL(−6) and (−2) induce weaker minor pauses ^46^. Pauses are a major determinant of transcription elongation kinetics as they last for a few seconds to minutes while the rate of elongation is typically in the range of 40 – 50 nt/sec ^22,23^.

Our results indicate that H2BK34ub facilitates transcription by suppressing pauses and shortening their durations at SHL(−6) and (−5) and that H2BK120ub also has the same effects but only to a moderate extent. These differential extents between H2BK34ub and K120ub are in good agreements with their differential efficiencies for Nap1-induced hexasome formation ^17^. It was only recent when H2BK34ub started getting attention while H2BK120ub has been actively investigated for a longer time for its role in recruiting Dot1L to methylate H3K79. The recent attention to H2BK34ub was triggered by the finding that H2BK34ub is coupled to H2BK120ub in a network of direct protein interactions. The interactions involve a general transcription elongation factor Paf1C and the two ubiquitin ligase complexes MOF-MSL and RNF20/40 for ubiquitylating H2BK34 and H2BK120, respectively, which eventually promote Pol II processivity *in vivo* ^16^. Since then, it has been shown that H2BK34ub nucleosomes have the ubiquitin moieties protrude between the two gyres of the nucleosomal DNA, weakening the structural integrity of the nucleosome and distorting DNA toward the entry region of the nucleosome ^20,21^. These results suggest that the effects of H2BK34ub on transcription kinetics can be the major mechanisms of facilitated transcription by H2B ubiquitylations. The effects are likely due to localized perturbations of the nucleosome structure near the DNA-(H2B-H2B) contact region, which has been validated by our recent structural study based on smFRET ^56^. We suspect that the weaker effects of H2BK120ub is due to its non-ideal location to affect DNA-histone interactions.

An important regulatory role for H2BK120ub is to recruit Dot1L and methylate H3K79. H3K79 methylations are enriched in active genes and its mis-regulation is a hallmark of many cancers ^57,58^. H3K79 methylations facilitate transcription *in vitro* and H3K79me_2_ alters the local histone surface without inducing any significant changes to the global structure of the nucleosome ^32,35,59,60^. The addition of methyl groups to H3K79 makes this location sterically bulky and can disrupt the weak hydrogen bonding between the lysine and the L2 loop of histone H4 ^35^. This change makes a hydrophobic pocket lined by H3L82 and H4V70 more solvent accessible ^35^, which may result in more flexible conformations around the H3K79 region. Such increased conformational flexibility would positively impact on the transcription efficiency. The shortened SHL(−2)/(−1) pause durations by H3K79me_3_ are in line with this mechanism. We suspect that the increased rate of elongation by H3K79me_3_ is likely due to shortened SHL(−2) pauses. As we included SHL(−2) pause durations in elongation time measurements, the shortened SHL(−2) pauses by H3K79me_3_ would result in an increased elongation rate. Overall, our results support that the increased flexibility around the (H3-H4)_2_ tetramer region facilitate transcription elongation by shortening the durations of SHL(−2)/(−1) pauses and subsequently elevating the rate of transcription elongation through this region, providing the physical mechanism of facilitated transcription by H3K79me_3_.

Another notable finding is that the effects exerted by H2BK120ub and H3K79me_3_ add up to facilitate transcription to the highest extent among all 5 nucleosomes that we examined (Fig. 6). According to our results and the previous studies, the mechanisms of these two modifications are localized in two different regions of the nucleosome – H2BK120ub near the SHL(−6)/(−5) region and H3K79me_3_ near the SHL(−2)/(−1) region. As such, the total effects would be the sum of their individual effects. Considering that H3K79me_3_ is induced by H2BK120ub recruiting Dot1L and these two modifications impact on the nucleosome structure in two different regions, we suggest that these two modifications may work together to contribute significantly to facilitated transcription *in vivo*. It is noteworthy that a previous research reported heightened nucleosome barriers around SHL(−2)/(−1) by H2BK120ub alone ^23^. Combined with our results, it is implied that the effect of H3K79me_3_ overwrites that of H2BK120ub around SHL(−2)/(−1). Considering that the end result of H2BK120ub is H3K79me_3_ upon transcription activation, we suggest that H2BK120ub helps maintaining the structure of the nucleosome around SHL(−2)/(−1) and induces H3K79me_3_ upon transcription activation to greatly facilitate transcription through the nucleosome.

A histone methyltransferase Dot1L mediates the crosstalk between H2BK120ub and H3K79me_3_ and is known to affect the structure of the nucleosome upon binding even in its inactive form ^28,29,61^. This effect of Dot1L might contribute to the mechanism of how Dot1L is coupled to facilitated chromatin remodeling ^62^. It will be an important future study to investigate how Dot1L binding to H2B ubiquitylated nucleosomes modulates the overall structure and stability of the nucleosome during transcription.

An important focus on the nucleosome dynamics during transcription is how the nucleosome maintains its structural integrity. A previous study suggested that H2A-H2B dimers are displaced or dissociated transiently and return to their original positions to reassemble the nucleosome during transcription elongation ^63^. Our results show that ∼30 % of the nucleosomes partially or fully rewrap the DNA around the entry proximal H2A-H2B contact region after Pol II passage and that H2BK34ub significantly facilitates partial rewrapping of DNA. This effect of H2BK34ub suggests its role for stabilizing nucleosome intermediates during transcription. The effect might be due to the favorable location of the ubiquitin to interact with DNA as stable interactions between ubiquitin and DNA have been suggested previously ^20,52,53^. Combining the effects of H2BK34ub, H2BK120ub, and H3K79me_3_ all of which are in the same regulatory network, our results suggest that these modifications induce and stabilize nucleosome intermediates during transcription. These dual roles would facilitate transcription elongation through the nucleosome while helping maintain the chromatin structure to ensure proper gene regulation.

## SUPPLEMENTARY INFORMATION

Supplementary Table S1, and Figures S1-S5.

## ACKNOWLEDGEMENTS

This research was funded by grants from NIH (R01GM123164 and R01GM130793 to T.-H.L.) and from IDB RAS government programs of basic research in 2022 № 0088-2021-0007 to W.K. We thank Dr. Joseph Reese for providing the yeast strain and plasmid for Pol II and TFIIS purification. We thank Dr. Song Tan for providing plasmids that encode human H2A and H2B histone.

