## Supplemental figures and tables for "The Effects of Histone H2B ubiquitylations and H3K79me_3_ on Transcription Elongation"

**Table S1**. Sequences of DNA oligos used to generate the transcription template containing Widom 601 nucleosome positioning sequence in our measurements.

**Figure S1**. Mass spectrometric analysis of H3K79 trimethylation.

**Figure S2**. SDS page analysis of the modified histone H2B.

**Figure S3**. Native PAGE analysis to confirm the presence of different sets of nucleosomes.

**Figure S4**. FRET distance estimation from Cryo-EM structures for two FRET pairs used in our measurements.

**Figure S5.** Sample FRET trajectories showing various times of transcription elongation.

**Table S1**. Sequences of DNA oligos used to generate the transcription template containing a Widom 601 nucleosome positioning sequence. "iAmMC6T" denotes the modification of thymine with C6 amine linker (Integrated DNA Technologies, Coralville, IA) for fluorophore labeling. The EC template region contains a 9-nucleotide mismatch that mimics a transcription bubble ahead of the nucleosome.

| DNA fragment | Sequence (5' – 3') |
|---|---|
| F1 | /5Phos/GCAG ATC GAG AAT CCC GGT GCC GAG GCC GCT CAA |
| F2 | /5Phos/ TTGG /iAmMC6T/CGT AGA CAG CTC TAG CAC CGC TTA AAC GCA CGT ACG CGC TGT CCC CCG CGT TTT AAC CGC CAA GGG GAT TAC |
| F2_alternate | /5Phos/TTG GTC GTA GAC AGC TCT AGC ACC GCT /iAmMC6T/ AAA CGC ACG TAC GCG CTG TCC CCC GCG TTT TAA CCG CCA AGG GGA TTA C |
| F3 | /5Phos/ TCC C/iAmMC6T/AGT CTC CAG GCA CGT GTC AGA TAT ATA CAT CCG AT |
| F3_alternate | /5Phos/TCC CTA GTC TCC AGG CAC GTG TCA GAT ATA /iAmMC6T/ ACA TCC GAT |
| F4 | /5Phos/GCG GCC GCG TAT AGG GTC CAT CAC ATA AGG GAT GAA CTC GGT GTG AAG AAT CAT GCT TTC CTT GGT CAT T/3Bio/ |
| R0 | AAT GAC CAA GGA AAG CAT GAT TCT TCA CAC CGA GTT CAT CCC TTA TGT GAT GGA CC |
| R1 | /5Phos/CTA TAC GCG GCC GCA TCG GAT GTA TAT ATC TGA CAC GTG CCT GGA GAC TAG GGA GTA ATC CCC TTG GCG GTT AAA ACG CGG GG |
| R2 | /5Phos/GAC AGC GCG TAC GTG CGT TTA AGC GGT GCT AGA GCT GTC TAC GAC CAA TTG AGC GGC CTC GGC ACC GGG ATT CTC GAT |
| EC Template | /5Phos/CTG CGC CAC CGC GGT CTA GAG GAT CCC CGG GAG TGG AAT GAG AAA TGA GTG TGA AGA GCT AAT TGA CTG ACG TAA GC |
| EC Non-Template | GCT TAC GTC AGT CTG GCC ATC TTT CAC ACT CAT TTC TCA TTC CAC TCC CGG GGA TCC TCT AGA CCG CGG TGG C |

**Figure S1**. Mass-spectrometric analysis confirms proper H3K79 trimethylation. The peak at 15301.4 m/z denotes the trimethylated H3 monomer (calculated molecular weight: 15299.7 Da)

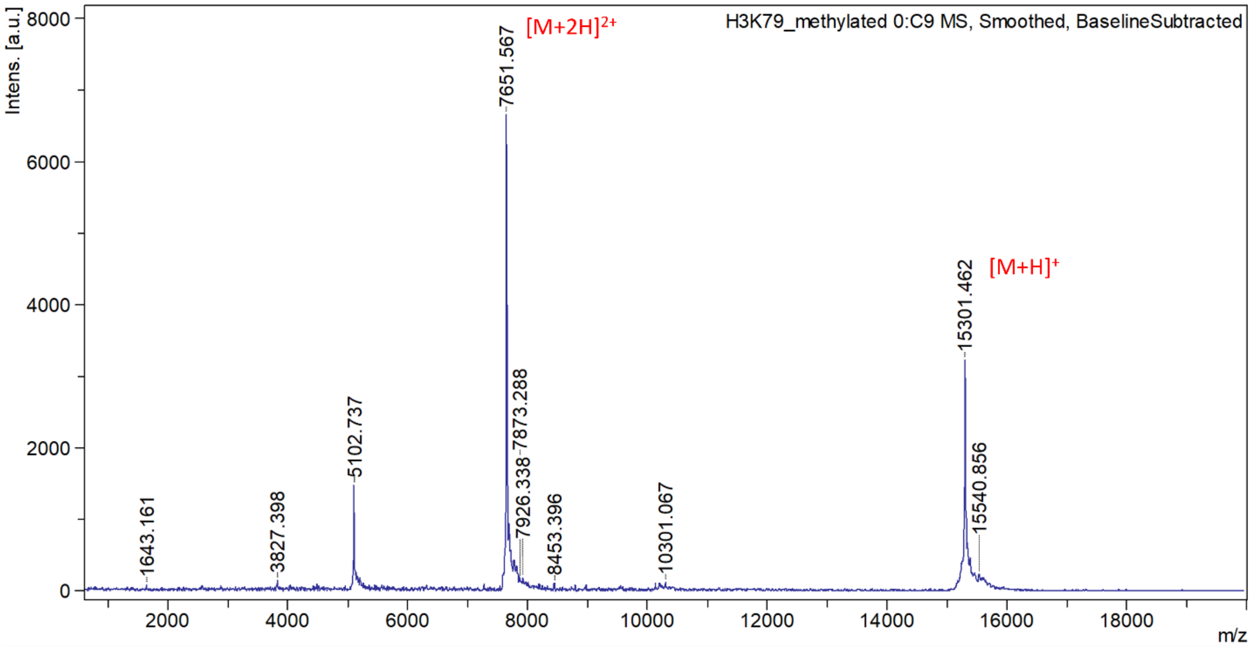

**Figure S2**. SDS page analysis confirms histone H2B ubiquitylations. The molar masses for H2B and ubiquitin are 14 kDa and 9 kDa, respectively.

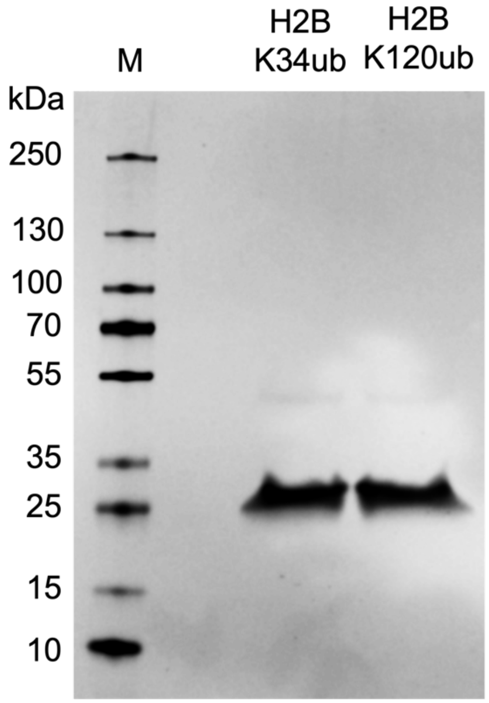

**Figure S3.** Native PAGE analyses confirm the assembly of the transcription templates containing the nucleosomes with the modifications marked on the top of the gel.

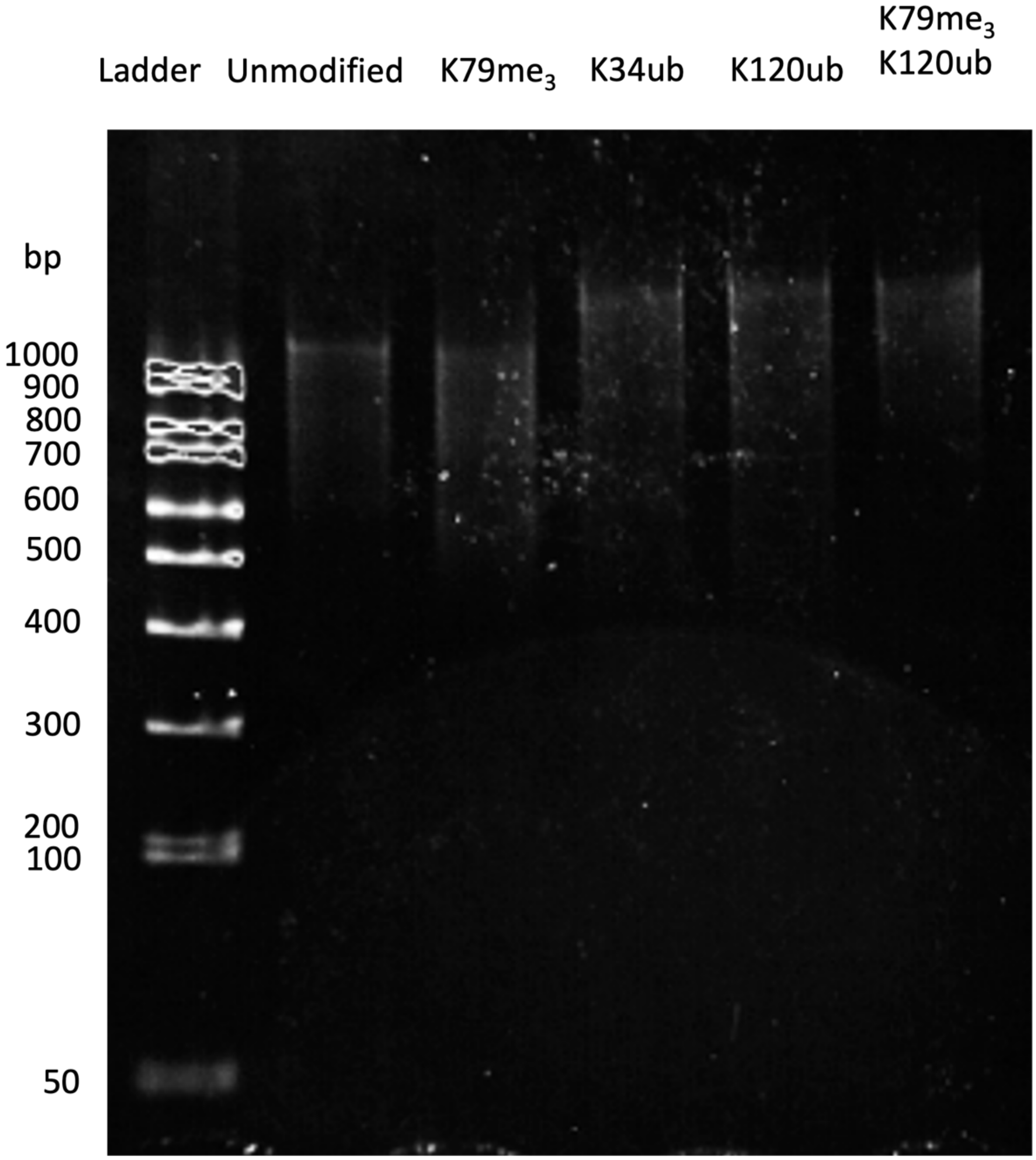

**Figure S4**. FRET distance estimation from Cryo-EM structures for two FRET pairs used in our measurements (Kujirai, T., Ehara, H., Fujino, Y., Shirouzu, M., Sekine, S.-i. and Kurumizaka, H. (2018) Structural basis of the nucleosome transition during RNA polymerase II passage. *Science*, **362**, 595-598). (A-D) The distances between the FRET donor and acceptor were estimated by measuring the distances between the two carbon atoms of the methyl groups at the 5'C atoms of the thymine bases at the +34$^{th}$ and +112$^{th}$ nucleotides in the pauses at SHL(-6), SHL(-5), SHL(2), and SHL(-1) (PDB 6A5O, 6A5P, 6A5R, 6A5T). (E-H) The distances between the FRET donor and acceptor in the alternate FRET pair (at the +57$^{th}$ and +138$^{th}$ nucleotides) were estimated using the same method.

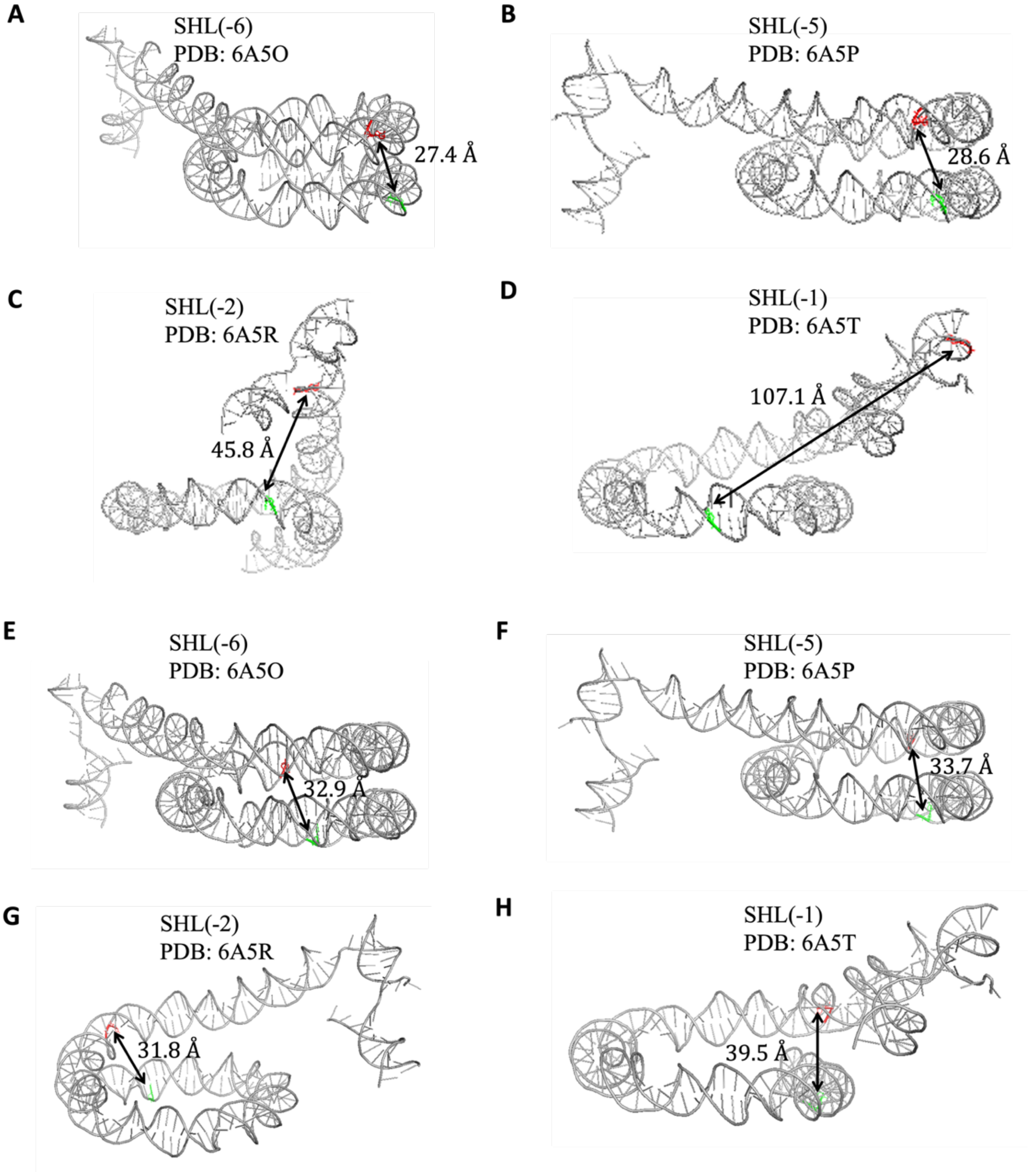

**Figure S5.** Sample FRET trajectories showing various times of transcription elongation. Green and red line denote Cy3 and Atto647N intensities, respectively. Blue lines represent the changes in the FRET efficiencies. (A-D) The elongation times are indicated by the arrows between the two dashed lines.

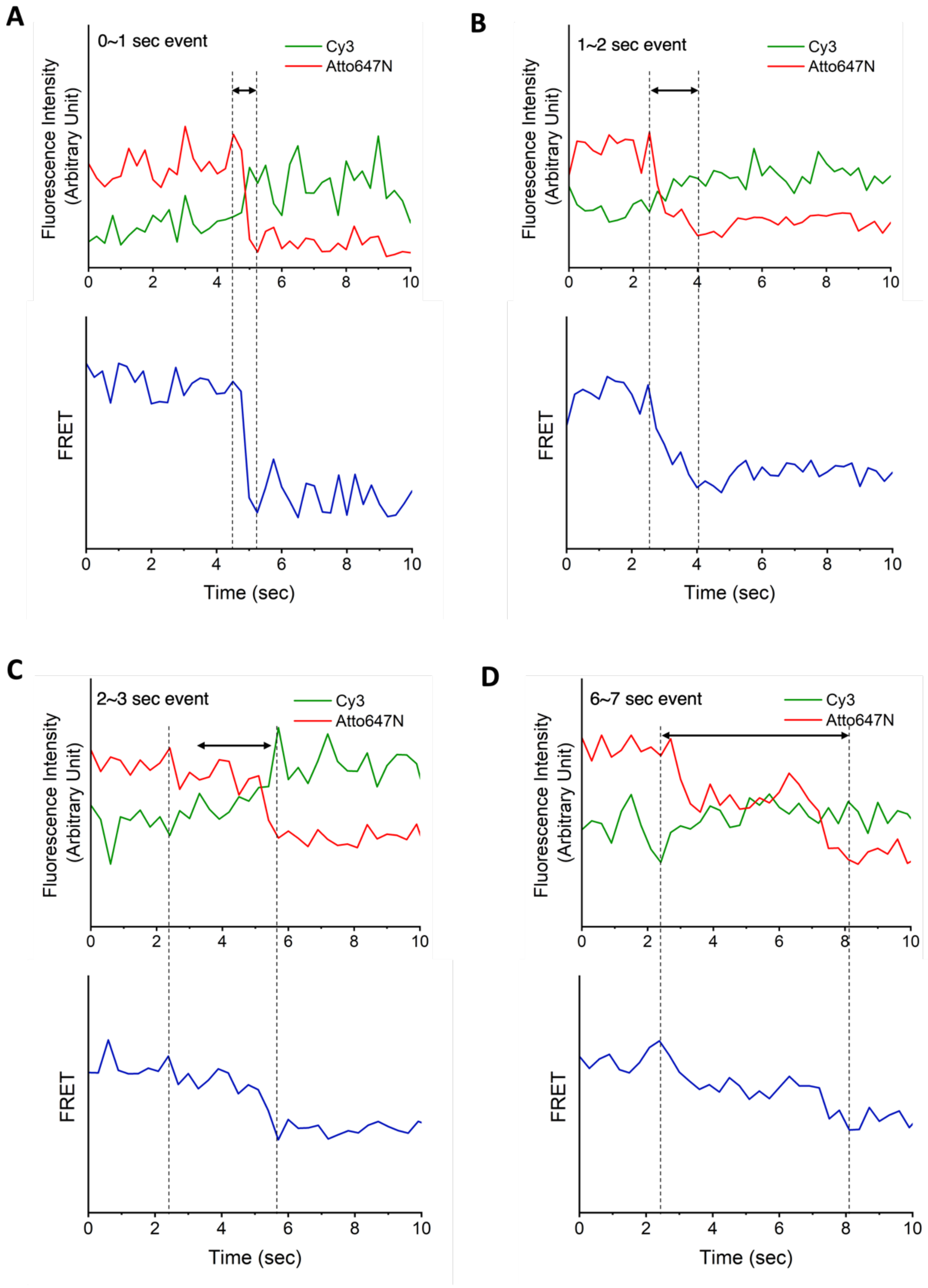
